# CNValidatron: Accurate And Efficient Validation of PennCNV Calls Using Computer Vision

**DOI:** 10.1101/2024.09.09.612035

**Authors:** Simone Montalbano, G. Bragi Walters, Gudbjorn F. Jonsson, Jesper R. Gådin, Thomas Werge, Daniel F. Gudbjartsson, Hreinn Stefansson, Andrés Ingason

## Abstract

**Background:** Large rare copy number variants (CNVs) are a main source of genetic variation in the genome and are important in both evolution and disease risk. CNVs can be detected using different data sources, including genome sequencing, genotyping arrays and quantitative PCR experiments, but in most large cohorts, genotyping arrays remain the most prevalent source. Current methods to call CNVs from genotyping array data suffer from high false positive rates and while multiple approaches, including QC filtering, visual inspection of intensity tracks, and wet-lab validation are commonly applied to counter this problem, such methods are often non-specific (QC filtering) or inefficient (visual and wet-lab validations) at a genome-wide scale.

**Results:** We have assembled the largest collection of human-verified CNV calls using visual validation, totaling almost 60,000 calls from 22,500 samples from three cohorts genotyped on several different arrays. Across all cohorts our visual validation found the majority of CNV calls to be false positive (53.7%) or unclear (9.7%). The false positive fraction varied substantially across datasets and genomic regions, and we show that existing filtering methods based on QC metrics are inefficient in removing false calls. Given the supremacy of visual validation over existing filtering methods in controlling the false positive fraction, we used a subset of our visual validation dataset to train a convolutional neural network to automate the validation of CNVs through machine vision. We tested the efficacy of the model using the remainder of the dataset and found the performance exceeded 90% in most measures, approximating that of a human analyst. Cross-validation with genome sequencing data found our visual validation to be highly accurate, with only 1.7% of calls supported by the sequencing dataset deemed as false by the human analyst, and a further 7.5% deemed as unclear.

**Conclusions:** Visual inspection is the only effective validation approach for CNV calls. Our model is capable of automating this task at scale with very high accuracy, as shown by testing both within-sample and out-of-sample. The software is available as an R package at https://github.com/SinomeM/CNValidatron_fl.

## Introduction

Copy number variants (CNVs) are a class of structural variants (SV), wherein segments of DNA have been deleted or duplicated resulting in an unbalanced chromosomal rearrangement^1,2^, although a specific definition to distinguish them from other structural variants does not exist in the literature. In this study we focussed on those deletions and duplications that can be detected from SNP array data, which most often are rare and relatively large. Therefore, for the purposes of this study we define CNVs as rare (<2.5% carrier frequency) deletions and duplications sized between small structural variants and large chromosomal rearrangements (ca. 50kb – 10Mb).

Whole genome sequencing (WGS) based on long-read technologies is currently considered the ideal data source for the identification of most types of SV, including CNVs, as it encompasses breakpoints of large and complex SVs within a single read^3^. However, such data is not yet available for large sample cohorts or biobanks. Existing large WGS datasets^4,5^ are mostly based on short-read sequencing, in which the presence of CNVs can be inferred from relative changes in read depth and allelic ratios, but only with limited accuracy^6–8^ whereby rare CNVs can be missed^9–11^.

CNVs can also be inferred from SNP array data through the same principles (i.e., relative change in DNA template and allelic ratios) and since SNP arrays remain the most prevalent source of human genotype data, such array-based data is still very relevant for the study of CNVs. For almost 20 years, PennCNV^12^ has been the most frequently applied method for SNP array-based CNV detection, especially in large sample collections. PennCNV uses the relative intensity of each SNP probe (defined as the log R ratio, LRR) and the ratio of the intensity of the two alleles measured for each probe (defined as the B allele frequency, BAF) to infer copy number state using a hidden Markov model (HMM).

The ability of PennCNV to detect true CNVs depends on factors such as sample quality, array type and genomic location, but it is generally considered to have a high sensitivity but poor specificity, with the false positive proportion of calls often exceeding 50%.^13^ To control this high false positive rate, researchers try to apply rigorous quality control, including refined filtering on various sample and CNV quality metrics^14^, or using the intersection of calls across different CNV calling algorithms^15,16^. However, a recent benchmarking study suggests that such cross-validation of calls, using different CNV-calling methods, does not improve upon the accuracy of PennCNV alone^17^, and the application of strict general filters may hurt the sensitivity to identify true CNVs.

In our experience, and that of many of our colleagues, the most reliable way to verify PennCNV calls from SNP array data (without external validation such as qPCR or WGS) is through visual inspection of the LRR and BAF tracks surrounding each call, as this allows for a multifaceted evaluation of the quality and context of the datapoints underlying the algorithm-generated call.^18–20^ However, this approach is very time consuming and therefore its application in large datasets has been limited to studies of relatively few genomic loci, known or suspected to harbor pathogenic CNVs.

In this study, we leverage CNV data from three different cohorts, involving multiple SNP arrays, a large sample of trios, and WGS-based CNV calls, to characterise the problem of efficient CNV validation from SNP array data, and present a solution to this problem based on machine vision. We showcase an algorithm that is able to automate the visual inspection of PennCNV calls, with an accuracy and precision comparable to that of a human analyst.

## Methods

### CNV calling and processing

This study is based on three main datasets: 15,000 randomly selected samples from the whole UK Biobank (UKB)^21^, 6,000 samples randomly selected from all individuals with full trio information (offspring with both parents genotyped as well) in the Icelandic biobank at Amgen deCODE genetics, and 1,500 samples from the 1000 Genomes study (1kG)^22,23^ each genotyped on two different SNP arrays and whole genome sequenced at high coverage. Samples were genotyped on different chips in different cohorts, Affymetrix UK Biobank Axiom Array in UKB^24^, multiple versions of the OmniExpress family of Illumina arrays in deCODE^25^, and both Illumina HumanOmni2.5 and Affymetrix Genome Wide 6.0 in 1kG. SNP maps were filtered to only include biallelic autosomal SNPs mapping uniquely to the Haplotype Reference Consortium (HRC) hg19 reference map^26^, with a minor allele frequency of at least 0.1%. All CNV calls were generated using PennCNV^12^. CNV calling was performed internally at deCODE for the Icelandic samples^20^, while we use our previously described protocol for the UKB and 1kG^27^. After first stitching adjacent PennCNV calls (spanning at least 5 markers) using the default settings of our previously published CNV protocol^27^, we excluded all calls that included fewer than 15 markers. Samples from deCODE had been pre-screened for QC outliers (standard deviation of LRR >0.3 and/or BAF drift >0.01). A schematic of the datasets used for the study is shown in figure 1.

**Figure 1.**
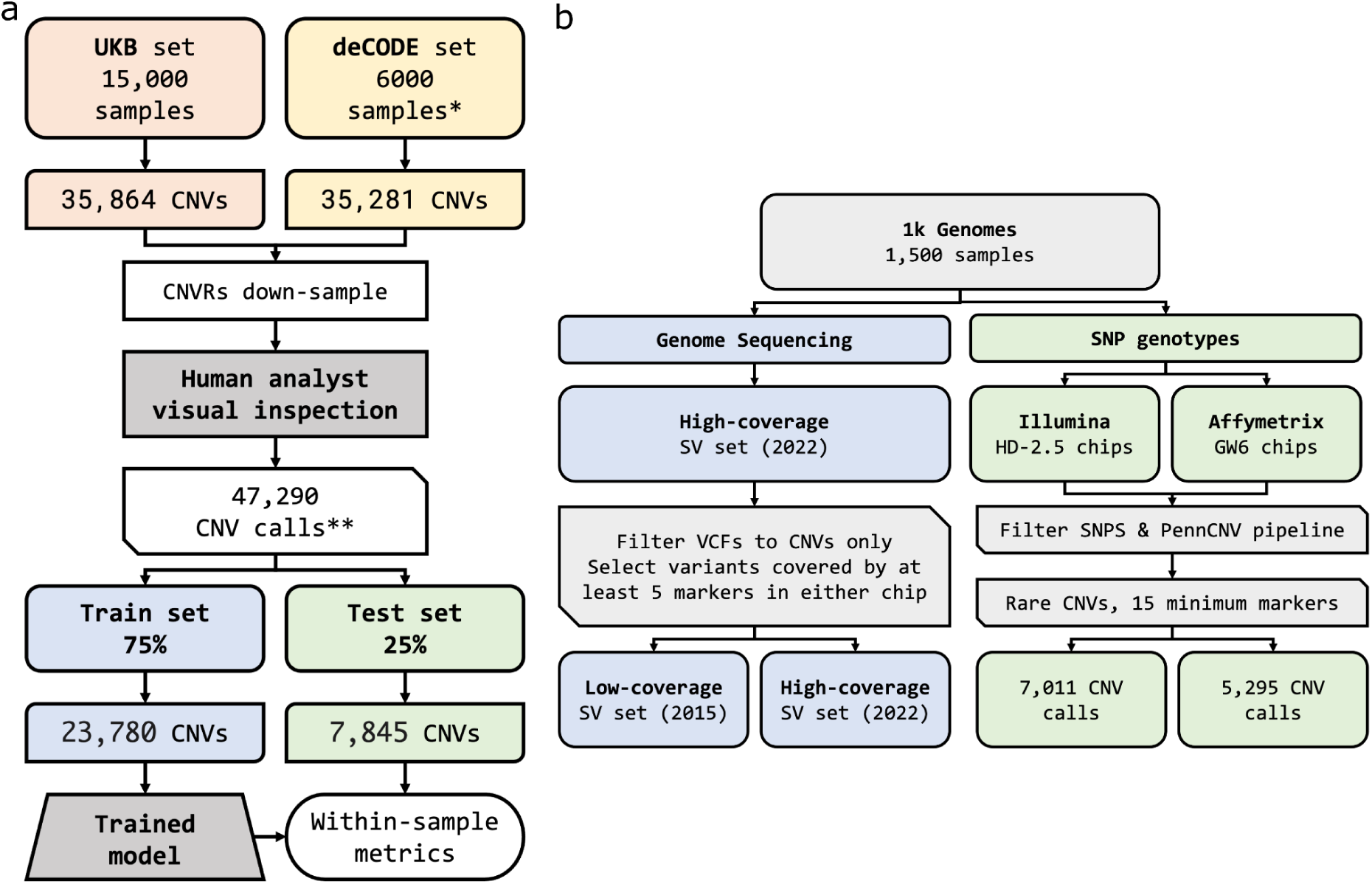
Schematic representations of the training and test sets and the overall processing pipelines. a) Train and within-sample test sets (UKB and deCODE). b) Out-of-sample test set (1kG). *: Samples with full trio information (both father and mother genotypes available) **: Including “Unclear”

All CNV calls were manually processed by a human analyst and assigned to one of three categories via visual validation: “True call” (deletion or duplication), “False call” and “Unclear call”. Visual validation refers to the manual inspection of the LRR and BAF patterns in the region of the putative CNV call in a given sample. Supplementary Figure 1 shows examples of different possibilities and a detailed explanation of the process. The visual validation was performed using an in-house graphical interface (https://github.com/SinomeM/shinyCNV).^27^

Since the distribution of CNV calls is not uniform across the genome, nor across different genotyping chips, for the training data (UKB and deCODE) we attempted to limit the number of examples from a specific locus. To do so we computed CNV Regions (CNVRs: homogeneous groups of CNVs with identical or near-identical breakpoints), and downsampled examples from the more common regions. This was done as follows: CNVRs containing up to 20 CNVs were not downsampled, CNVRs with 21 to 50 CNVs were downsampled to 20 CNVs, CNVRs with 51 to 100 CNVs were downsampled to only 30 CNVs and CNVRs with more than 100 CNVs were downsampled to 40 CNVs. After this step, the number of CNVs in UKB and deCODE was 23,056 (15,609 deletions and 7,447 duplications) and 24,234 (17,287 deletions and 6,947 duplications) respectively. The algorithm to compute CNVRs is described in detail below.

### CNVR detection algorithm

The CNVR detection algorithm is composed of three separate steps. First, CNVs are divided into networks, defined as the smallest set of mutually overlapping CNVs. By definition, the largest possible network is thus a chromosomal arm, and the smallest network is a single CNV without overlap with any neighbour. In our pipeline CNVs cannot span across centromeres, due to lack of marker coverage.

The second step is where the CNVRs are created. For each network, an *N* × *N* matrix is constructed, where *N* is the number of CNVs in the networks; each *xy* value is the intersection 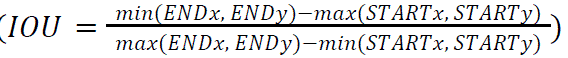 CNVs will have an IOU of 1, two overlapping CNVs will have an IOU between 0 and 1 depending on the similarity, and two non-overlapping CNVs will have an IOU closer to –1 the more distant they are. This matrix is then translated into a weighted igraph^28^ network object where all pairs with an IOU above a set threshold (0.75 in this study) will be connected, with the IOU values being the strength of the connection. This network is then processed with the community detection algorithm Leiden^29^ (igraph::cluster_leiden(), with the resolution set to 1), and the resulting networks are converted back into CNVRs with boundaries set at median values of the start and end of constitutive CNVs, respectively. CNVs that do not cluster with any other calls (i.e., singletons) are considered their own CNVR.

The third and final step is an optional forced merge of CNVRs. For each chromosome, all CNVRs are processed in genomic order by start position. For any given CNVR, all regions with an IOU above a certain threshold (default value is 0.75) are merged together and the new start and end are computed again as the median from all the CNVs in the new group. This process continues until all CNVRs have been either processed or merged with a previous one. This is done for a fixed number of iterations, 5 by default.

### One Thousand Genomes Analysis

For genotyping, two different chips were used, Illumina HD 2.5 (referred to as HD hereafter) and Affymetrix Genome Wide 6 (referred to as GW hereafter). Intensity data was processed in the same way as the UKB, and CNVs were called using the same PennCNV pipeline and parameters. Importantly, these sets differed substantially from UKB and deCODE in both the density of the genotyping chips (∼705,000 markers in HD and ∼900,000 in GW after filtering), as well as the actual method to obtain the sample (blood vs cell lines). CNVR-downsampling was deemed unnecessary in this set given the lower sample count and denser arrays.

In addition to SNP array data, we obtained a dataset of structural variants that had been called from high-coverage (targeted read-depth 30x) genome sequencing of the 1kG samples^30^. The structural variants were obtained as VCFs and loaded in R using the package VariantAnnotation^31^. Then we selected only variants marked as “CNV” and further defined as “detectable” those covered by at least 5 markers in either array to constitute the final datasets.

When subsequently comparing the agreement between any two CNV sets, A and B, from the three available sources for the 1kG data (the two different SNP arrays and the genome sequencing), we considered a segment in set A “replicated” in B when a segment with the same dosage change was found to have an IOU of at least 0.3 in set B.

### Prediction model design and training

The model was trained on a combined set of UKB and deCODE samples, consisting of 75% of the total set for both (11,250 and 4,500 samples, respectively). In these training data, very ambiguous calls (i.e., those still marked as “Unclear” after two inspections and/or comparison with LRR/BAF tracts of parents) were excluded (Supplementary Table 1). Training was performed both on the two sets separately (UKB, 11,250 examples, or deCODE, 12,530 examples) and on the combination of the two (23,780 examples). As part of the training pipeline, we subsequently increased the number of training examples using data augmentation in the form of image flipping on the x axis with a probability of 0.5 per example.

The training model was built using torch^32^ with the R packages torch^33^, torchvision^34^ and luz^35^. It is composed of 5 convolutional layers (nn_conv2d) followed by a 10-layer linear classifier (nn_linear). Each layer has a 5% dropout rate (nn_dropout2d(p = 0.05). Training was done using luz::fit() with the following settings: maximum learning rate of 0.02, maximum epochs of 50, and early stopping with a patience value of 4. The training set was further divided into a 75-25% configuration for training and validation subsets as suggested by the luz and R torch developers.

The data representation was designed to be a simplified and condensed version of what the human analyst sees when performing the visual inspection of CNVs (supplementary figure 1). All the processing performed has two main objectives, first to fit all necessary information in the smallest amount of data possible (here, the smallest picture) to minimise computation, and second to make the pictures as similar as possible across different genomic loci, different CNVs sizes, and different number and density of SNP markers in the region of the CNV call. We loosely tested different resolutions before setting it to 96 by 96 pixels. As shown in supplementary figure 1, the picture is composed of three rows, containing two views of the LRR values around the CNVs and one view of the BAF. Horizontally the image is composed of three sectors, markers on the right of the CNV call, markers within the CNV call and finally on the left of the CNV call. Depending on the size of the CNV call (below 100kbp, between 100kbp and 1Mbp, above 1Mbp) the two bottom rows will cover 19, 15 or 11 times the length of the putative CNV, while the top row is always zoomed out by a further factor of three. The pixel image is constructed as follows. For each of the three rows, all points are moved to the coordinate system, i.e., from genomic position in x and LRR/BAF in y to 0-96 in x and [a,b] in y (where a and b values depend on the specific row). Then, the intensity of each pixel is computed as the logarithm of the number of points at the coordinate, scaled to the interval [0,1]. To minimise the effect of differential SNPs density across the locus of interest this process is performed in windows. Window size is 4, 8 or 32 pixels, depending on the CNV length class, as described above.

### Testing the model performance

The model performance was then tested in the 25% of UKB and deCODE samples that had not been used for the model training (within-sample test set), as well as out-of-sample in the 1kG set. In the within-sample test set “Unclear” calls were excluded, whereas in the out-of-sample test set (1kG) they were treated as “False”. In the context of the model performance we consider as; true positives (TP) those calls predicted true by the model with 0.5 probability or more that were evaluated as “True” by the human analyst; false positives (FP) those calls predicted true from the model (probability >= 0.5) but not evaluated as “True”; true negatives (TN) those calls evaluated as “False” and having a probability below 0.5; and false negatives (FN) those calls evaluated as “True” but having a probability below 0.5. In order to test the model performance we used accuracy, the F1 score, and the AUC (area under the curve). Accuracy is defined as 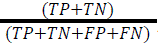 F1 score defined as 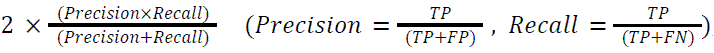, and the AUC was computed using the R package pROC.

### Software availability

The software described is available as an R package at https://github.com/SinomeM/CNValidatron_fl.

The pretrained model can be downloaded separately at https://doi.org/10.5281/zenodo.17174637.

## Results

### CNV calling results and validation

We called CNVs using a PennCNV pipeline^27^ in four datasets, identifying 35,864 (22,270 deletions, 13,144 duplications), 35,281 (23,147 deletions, 12,134 duplications), 15,866 (10,897 deletions, 4,969 duplications) and 11,916 (6,256 deletions, 5,660 duplications) CNVs for UKB, deCODE, 1kG HD, and 1kG GW, respectively. As detailed in the Methods, in UKB and deCODE we subsampled CNVRs with more than 20 carriers, while in 1kG we excluded CNVRs with a frequency above 2.5%. This resulted in 23,056 (15,609 deletions, 7,447 duplications), 24,234 (17,287 deletions, 6,947 duplications), 7,011 (4,898 deletions, 2,113 duplications) and 5,295 (2,879 deletions, 2,416 duplications) calls in UKB, deCODE, 1kG HD, and 1kG GW respectively.

All remaining CNV calls were then processed by at least one human analyst through visual evaluation and marked as either “True”, “False” or “Unclear” (Supplementary Figure 1 and Supplementary Table 1). We observed a similar distribution of true and false positives across UKB, deCODE and the 1kG-HD datasets, with more than 50% of the total calls being false, whereas in the 1kG-GW we deemed only ∼20% of calls to be false positives.

### Model training and testing

We used the human-validated CNV calls from 75% of samples from UKB and deCODE to train a convolutional neural network to discriminate between the three following categories: “False call”, “True Deletion” and “True Duplication”. Training for the joint model stopped after 13 epochs with an accuracy of 0.96.

We plotted the precision and recall in the within-sample test set (i.e., testing the performance of the model in predicting calls in those samples from UKB and deCODE that had not been used in training the model) across all probability cutoffs (Figure 2) and selected 0.5 as a default value to base our performance evaluations. As shown in Table 1, precision, recall, accuracy, F1 score and AUC are all well above 0.9 both within-sample (UKB and deCODE test data) and out-of-sample (1kG HD and GW data), with the exception of the accuracy in the 1kG-GW set at 0.89.

**Figure 2:**
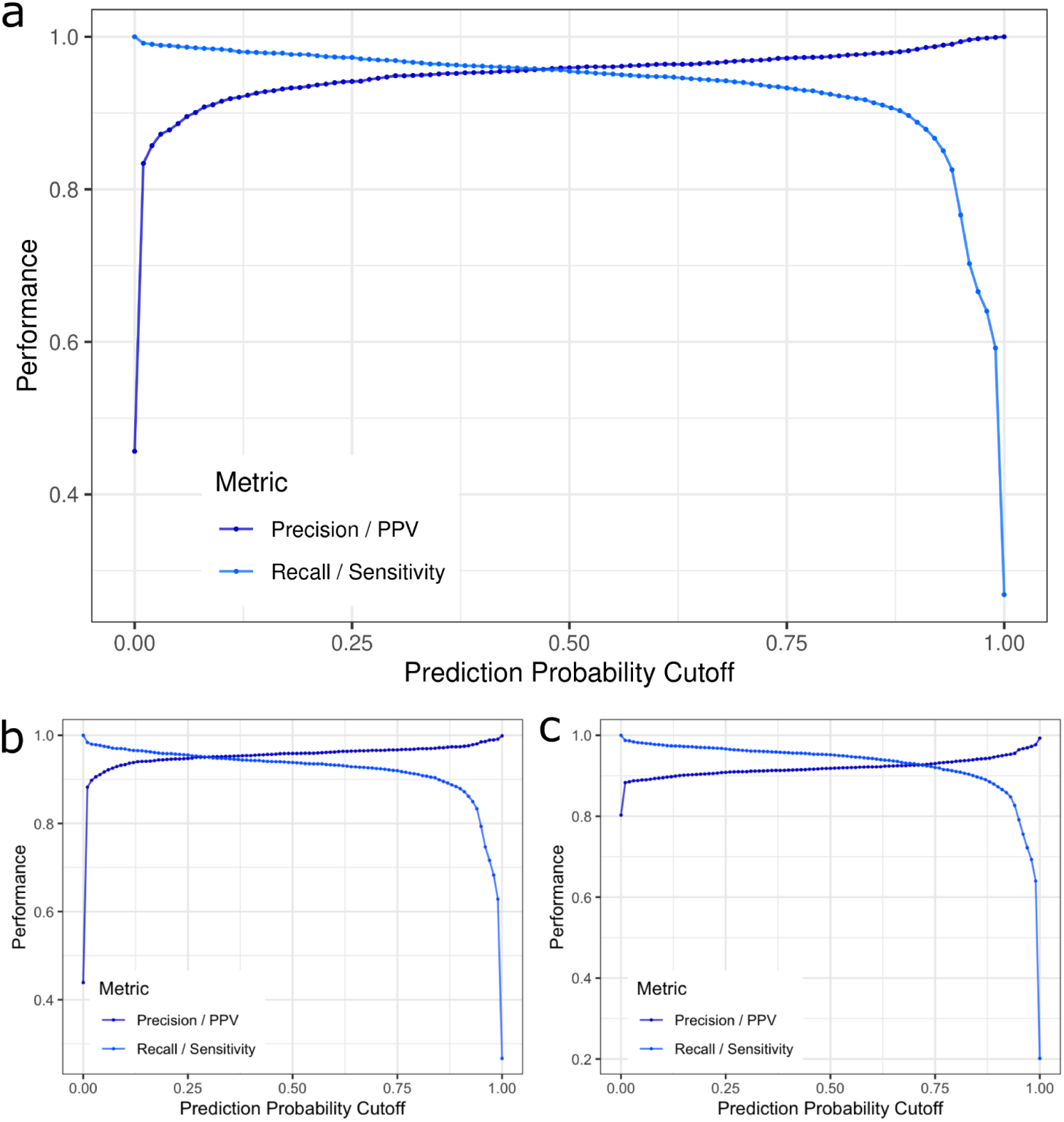
Performance metrics (precision or positive predictive value in dark blue and recall or sensitivity in light blue) of the model over the prediction probability. a) within-sample combined test set (25% of UKB + deCODE), b) 1kG Illumina HD, c) 1kG Affymetrix GW. “Unclear” calls are excluded from the training and within-sample test set (a), but are treated as “False” in the out-of-sample test sets (b-c).

**Table 1:** Model performance metrics at 0.5 prediction probability cutoff in all available test sets, within-sample (UKB and deCODE samples that were not used for model training) and out-of-sample (1kG HD and GW). TP = true positive, TN = true negative, FP = false positive, FN = false negative, AUC = area under the curve.

| Testset | TP | TN | FP | FN | Precision | Recall | Accuracy | F1 score | AUC |
| --- | --- | --- | --- | --- | --- | --- | --- | --- | --- |
| 1kG (HD) | 2888 | 3809 | 124 | 190 | 0.959 | 0.938 | 0.955 | 0.948 | 0.987 |
| 1kG (GW) | 4047 | 684 | 360 | 204 | 0.918 | 0.952 | 0.893 | 0.935 | 0.916 |
| UKB | 1656 | 1832 | 92 | 89 | 0.947 | 0.949 | 0.951 | 0.948 | 0.989 |
| deCODE | 1760 | 2290 | 53 | 73 | 0.971 | 0.960 | 0.970 | 0.965 | 0.993 |

We also tested whether a model trained only in one sample would perform better; thus we trained two additional models using UKB examples only or deCODE examples only. As expected, the models trained in one cohort performed better within cohort than across (i.e., models trained on UKB resulted in higher accuracy in UKB than in deCODE and vice versa). However, the model trained on the joint set was overall better (Supplementary Table 3).

### Validating the human visual inspection

In the 1kG data, we used the set of structural variant calls obtained from high-coverage sequencing^30^ as an orthogonal validation of our visual inspection in genotyping chip data. Given that the different call sets are obtained from completely independent sources, we assume that a CNV present in both array and sequencing data is very likely to be a true positive. For the 1kG-HD set, we found 2,576 calls to be replicated in the SV set, of which 2,430 (94.3%) were evaluated as “True”, and 114 (5.6%) as “Unclear” by the human analyst. For the 1kG-GW set, we found 3,936 calls to be replicated in the SV set, of which 3,487 (88.6%) were evaluated as “True”, and 372 (9.5%) as “Unclear” by the human analyst. Overall, less than 2% of these sequencing-supported PennCNV calls were marked as “False” by a human analyst.

Notably, 502 (16.3%) in 1kG-HD, and 764 (18.0%) in 1kG-GW “True” CNV calls from array data were not supported in the sequencing variants. Finally, we also compared the CNV calls from the two different arrays (HD and GW) with each other. We found 2,914 replicated in both sets. Of these, only 37 (1.3%) were “discordant” meaning they were evaluated as “True” in one set and “False” in the other. Taken together these results suggest that our visual validation is very accurate and also consistent across different analysts and genotyping platforms.

### PennCNV accuracy and QC metrics

Based on our experience and the previous literature consensus, we expected a link between the CNV validation (“True” / “False” / “Unclear”) and metrics regarding the specific call (number of SNP markers and call length) as well as of the sample QC metrics (LRR standard deviation (LRRSD), BAF drift and GC waviness factor (GCWF)). In UKB and deCODE data, true positives tend to be larger than false positives (all p values < 0.005, Student’s t test) and to include more markers (all p values < 2e-16, Student’s t test), supplementary figure 3a-d. Moreover, sample-wise quality metrics also have an impact on PennCNV accuracy (supplementary figure 4 a-d), with false positives tending to have higher LRRSD and BAF drift (all p values < 2e-16, Student’s t test). We found true positive deletions to be on average shorter and called from fewer markers than true positive duplications (all p values < 2e-16, Student’s t test). Finally, we pondered whether true and false positives follow the same distribution across the genome or not. As shown in supplementary figure 5-8, there is a mixture of patterns. While in some parts of the genome both true and false positives are present, others are mainly characterized by one category only.

In 1kG data, true positives had a higher number of SNPs in the GW set (p value < 2e-16, Student’s t test) but not in the HD set, supplementary figure 3 e-h. However, true and false positives did not differ in average length in either set (all p values > 0.5, Student’s t test). Within true positives, duplications were larger and included more markers in both sets (all p values < 0.0003, Student’s t test). Regarding QC measures (supplementary figure 4 e-h), true and false positives significantly differed in LRR SD, BAF drift and absolute WF in both sets (all p values < 2e-16, Student’s t test), while no significant difference was observed between deletions and duplication within the true positives (all p values > 0.2, Student’s t test). Finally, despite the smaller size of the CNVs sets, we also observe similar patterns to UKB and deCODE in terms of differential distribution of true and false positive calls (supplementary figure 9-12).

### QC filtering strategies

Based on the significant differences across QC metrics between true and false positives calls, we tested how standard filtering procedures would perform in reducing the false positives burden in the UKB and 1kG set (data from deCODE samples was obtained pre-filtered). For UKB we used the same filter as in Kendall et al.^36^, excluding samples with a |WF| > 0.03, more than 30 CNVs or a LRRSD > 0.35. This excluded 788 samples (5.3%) accounting for 204 true (2.8%), 3465 false (30.8%) and 168 unclear (3.7%) calls. While this filtering retains almost all the true positives, it only removes around one third of the false positives. In the 1kG set we used a similar “naive” QC-filtering approach^27^, excluding samples with a LRRSD > 0.35 a BAF drift > 0.01 and a |WF| > 0.03. This excluded 786 samples (52.4%) accounting for 1,564 true (50.8%), 3,218 false (89.0%) and 235 unclear (77.0%) calls in the 1kG-HD set, and 49 samples (3.3%) accounting for 107 (2.5%) true, 400 false (83.5%) and 51 unclear (10.6%) calls in the 1kG-GW set. In contrast to the UKB dataset, here the filtering strategy was very effective in removing false positives in both arrays. However, in the 1kG-HD set it unexpectedly also excluded more than half of the samples. We found this to be due almost exclusively to the WF filter.

Finally, we tested a more refined filtering method developed by Macé et al.^14^. Notably, we found the recommended filtering thresholds of 0.8 and 0.5 to be overly aggressive in all three sets, and we obtained more accurate results using a much lower threshold value of 0.2 (Supplementary Table 2). The filtering algorithm was able to remove the majority of the false (>90%) and unclear calls (>70%), however, it also excluded from 30% to >50% of the true positives (Supplementary Table 2).

## Discussion

In this study we created the largest collection of systematically validated CNV calls from genotyping data, consisting of data from three different cohorts (UK Biobank, Icelandic biobank at deCODE genetics, and the One Thousand Genomes Project). The samples we included were genotyped on multiple arrays from two major manufacturers (Illumina and Axiom Affymetrix/Thermo Fisher) and notably, all included 1kG individuals were genotyped on two distinct arrays. Although newer and potentially more precise methods to detect CNVs exist, genotype data still represent an important data source for research, such as association studies in large human collections.

Overall, we found more than half of the CNV calls produced by PennCNV (the most widely used CNV calling method for SNP arrays) to be false positives, although in the 1kG set genotyped on the Affymetrix Genome Wide 6 chip only 20% of calls were deemed false or unclear. While it is impossible to rule out differences in sample handling and processing, this could suggest that some arrays are better suited than others for CNV calling. Notably, we do not observe strong differences in the proportion of true CNVs and false positives between UKB (newer Axiom arrays) and deCODE (several subtypes of Illumina arrays). Finally, as the CNV validation in our study is based on human judgement, we use CNV calls from a completely independent structural variants pipeline based on high coverage whole genome sequencing data in the 1kG samples to demonstrate the accuracy of our evaluations.

Consistent with our expectations, we see that true CNV calls tend to be larger and include more markers than false positives in UKB and deCODE data. However, this pattern is much less clear in 1kG, which have been genotyped on denser arrays, suggesting that false positives might be partially correlated to marker density. True duplication calls tend to be larger and to include more markers than deletions across all the datasets. This is again in line with our experience and expectations, as the relative change in LRR is less for duplications than deletions, and their validation thus relies to a larger degree on a sufficient number of heterozygous genotypes to establish a clear BAF pattern typical of duplications. False positive calls were also more likely to belong to samples with higher LRRSD, BAF drift and absolute WF. This is of course widely recognised; in fact, most CNV studies begin with a QC-filtering step to remove “bad” samples.

We used this large dataset to train a CNN with the goal of automating the human visual inspection, reasoning that since the manual approach works well, while failing to scale properly, it would be the perfect task for a machine vision solution. Using extensive validation data, both within-sample and out-of-sample, we show that the model has a very high accuracy, and performs especially well in data sets with a large proportion of false positive calls. Despite the good overall performance and the large range of chips used in this study, we still recognise that every genotyping chip and, perhaps more importantly, each large genotyping cohort, is different. For this reason we recommend creating a small validation set when using the tool in a new dataset in order to gauge, and possibly calibrate, the cohort-specific accuracy. In our view, this marks an important improvement in automated filtering approaches to array-based CNV data, as we find that both naive^27,36^ and refined^14^ existing filtering strategies are at best only partially successful in discarding false positive calls, and in some cases sacrifice a large proportion of samples in the filtering process.

When plotting the genomic distribution of validated CNV datasets we observe some clear differences between the distribution of true calls and false positives. Thus, in some genomic regions most calls represent true CNVs, while in other regions most calls are false positives. While some of these hotspots of false calls tend to aggregate close to the telomeric regions of some chromosomes across most array types, other hotspots seem to be largely array-type specific. Studies that do not use a strong method to exclude false positive calls (such as visual inspection) are therefore at risk of producing highly biased results, especially when not taking account of array type when analysing samples with genotypes from multiple arrays.

The main limitation of this study is that we use only one algorithm, PennCNV, to generate the initial CNV dataset, while other alternatives were also available. However, we do not think this is a strong limitation as, 1) PennCNV is by far the most used software to call CNVs from genotype data, and 2) most programs (with the notable exception of ipattern^37^) use both the same raw data (LRR and BAF) and some implementation of the same HMM architecture as PennCNV.^38,39^ Also, a recent study that compared different CNV calling methods found that no other single method, or cross-validation across different methods, increased the (limited) accuracy achieved by using PennCNV alone.^17^

In conclusion, CNValidatron is a neural network which we have meticulously trained to distinguish between true CNVs and false positive calls in PennCNV-derived CNV call sets across different SNP arrays with high accuracy. It is our hope that the use of this tool, and continued improvement on it, will increase the scope, efficiency, and quality of CNV studies in large datasets for years to come.

## Declarations

### Ethics approval and consent to participate

The use of Icelandic data was approved by the National Bioethics Committee (NBC) (nos. VSN-16-067 and VSN-21-24). All genotyped participants signed a written informed consent allowing the use of their samples and data in projects at deCODE genetics approved by the NBC. Data was anonymised and encrypted using a third-party system, approved and monitored by the Icelandic Data Protection Authority.^40^

The UKB data were obtained under application no. 42256. All participants provided written informed consent for the use of genotype data and link to electronic health records. The North West Research Ethics Committee reviewed and approved the UKB protocol (ref. 06/MRE08/65). The 1000 Genomes Project data were obtained from The International Genome Sample Resource (internationalgenome.org) and are consented for public distribution without access or use restrictions.

### Consent for publication

Not applicable.

### Availability of data and materials

The code is available on GitHub https://github.com/SinomeM/CNValidatron_fl. The repository also includes the CNV and SV tables of the 1kG samples. The trained model is available on Zenodo at https://doi.org/10.5281/zenodo.17174637. UKB data is available to affiliated researchers through the official channels https://www.ukbiobank.ac.uk/. The Icelandic data at deCODE is not publicly available.

### Authors’ contributions

SM and AI designed the study, performed the visual validations and wrote the first draft of the manuscript; SM, GBW, GFJ, and JRG gathered and preprocessed array intensity data; SM developed the software and trained the CNValidatron model; TW, DFG, HS and AI helped secure access to the data and funding of the study; all authors contributed to the final version of the manuscript.

## Acknowledgements

We thank all participants of the three studies for making this study possible, among many others. We also thank the 1000 Genomes project for the data availability.

## Supplementary material

**Supplementary Table 1.** Human visual inspection results (true/false/unclear) in the different sets, divided by CNV genotype (deletion/duplication). UKB (15,000 samples), deCODE (6,000 samples), 1kG (1,500 samples). In 1kG, results for the two arrays are reported separately. The percentage of the total of each group is reported in parenthesis.

| Human Evaluation | CNV genotype | N UKB (%) | N deCODE (%) | N 1kG HD (%) | N 1kG GW (%) |
| --- | --- | --- | --- | --- | --- |
| True | Deletions | 4262 (18.5%) | 3194 (13.2%) | 1466 (21.0%) | 2195 (41.5%) |
| True | Duplications | 2959 (12.8%) | 4047 (16.7%) | 1612 (23.0%) | 2056 (38.8%) |
| False | Deletions | 7886 (34.2%) | 13863 (57.2%) | 3256 (46.4%) | 473 (8.9%) |
| False | Duplications | 3356 (14.6%) | 2733 (11.3%) | 372 (5.3%) | 92 (1.7%) |
| Unclear | Deletions | 3461 (15%) | 230 (0.9%) | 129 (2.5%) | 211 (4.0%) |
| Unclear | Duplications | 1132 (4.9%) | 167 (0.7%) | 176 (1.8%) | 268 (5.1%) |

**Supplementary Table 2.**
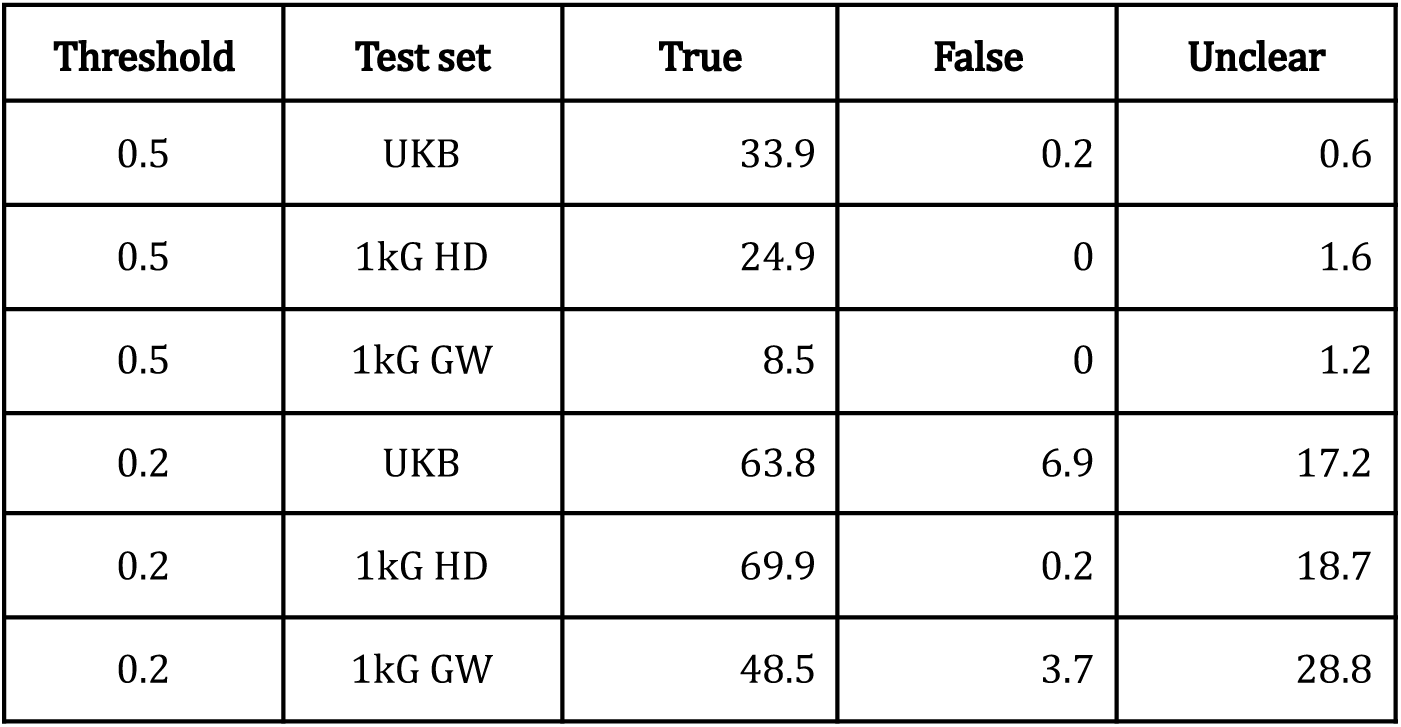
Percentage of calls in the three categories (True/False/Unclear) retained after applying the QC cleaning algorithm proposed by Macé et al.^14^ at two different thresholds, 0.5 (strict) and 0.2 (loose).

| Threshold | Test set | True | False | Unclear |
| --- | --- | --- | --- | --- |
| 0.5 | UKB | 33.9 | 0.2 | 0.6 |
| 0.5 | 1kG HD | 24.9 | 0 | 1.6 |
| 0.5 | 1kG GW | 8.5 | 0 | 1.2 |
| 0.2 | UKB | 63.8 | 6.9 | 17.2 |
| 0.2 | 1kG HD | 69.9 | 0.2 | 18.7 |
| 0.2 | 1kG GW | 48.5 | 3.7 | 28.8 |

**Supplementary Table 3.** Performance of the three models (trained on UKB data only, deCODE data only or both) in the three possible within-sample test sets (UKB only, deCODE only and both).

| Model | Test set | TP | TN | FP | FN | Precision | Recall | Accuracy | F1 score | AUC |
| --- | --- | --- | --- | --- | --- | --- | --- | --- | --- | --- |
| joint | joint | 3416 | 4122 | 145 | 162 | 0.959 | 0.955 | 0.961 | 0.957 | 0.992 |
| joint | ukb | 1656 | 1832 | 92 | 89 | 0.947 | 0.949 | 0.951 | 0.948 | 0.989 |
| joint | deCODE | 1760 | 2290 | 53 | 73 | 0.971 | 0.960 | 0.970 | 0.966 | 0.993 |
| ukb | joint | 3164 | 4061 | 206 | 414 | 0.939 | 0.884 | 0.921 | 0.911 | 0.972 |
| **ukb** | **ukb** | 1677 | 1794 | 130 | 68 | 0.928 | 0.961 | 0.946 | 0.944 | 0.983 |
| ukb | deCODE | 1487 | 2267 | 76 | 346 | 0.951 | 0.811 | 0.899 | 0.876 | 0.963 |
| deCODE | joint | 3279 | 3959 | 308 | 299 | 0.914 | 0.916 | 0.923 | 0.915 | 0.976 |
| deCODE | ukb | 1528 | 1666 | 258 | 217 | 0.856 | 0.876 | 0.871 | 0.865 | 0.942 |
| **deCODE** | **deCODE** | 1751 | 2293 | 50 | 82 | 0.972 | 0.955 | 0.968 | 0.964 | 0.994 |

**Supplementary figure 1.**
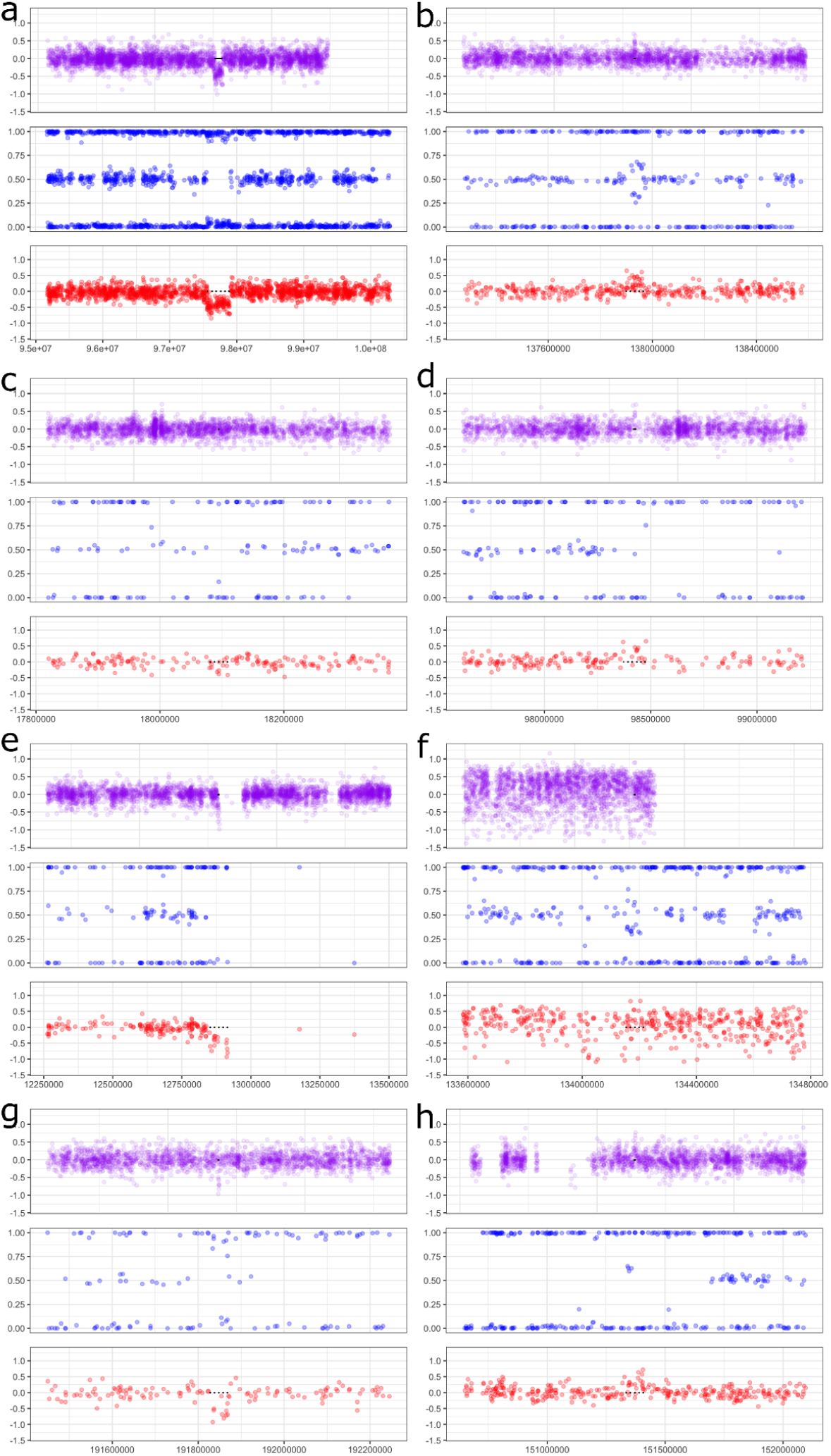
Examples of True (a-b), False (c-d) and Unclear (e-h) CNV deletions (a, c, e, g) and duplications (b, d, f, h). The characteristic patterns (lower LRR and LOH in the BAF for deletions, elevated LRR and split of the central BAF band for duplications) are very clear in the “true” examples and absent in the “false” ones. The unclear, as the name suggests, are less obvious. For deletions e) shows a centromeric region where LOH is present but the LRR decreases in a wave rather than sharply. This could be compatible with both noise (genomic waviness) and a true but not “clean” deletion. In g) another poor deletion is shown, here a larger LOH region combined with noisy LRR makes it harder to unequivocally confirm the call. For duplications, example f) shows some indication of a true duplication in the BAF but the LRR is very noisy and not elevated cleanly enough to confirm it. Finally, in h) we see a larger region of LOH that prevents the validation of the somewhat elevated LRR signal.

**Supplementary figure 2.**
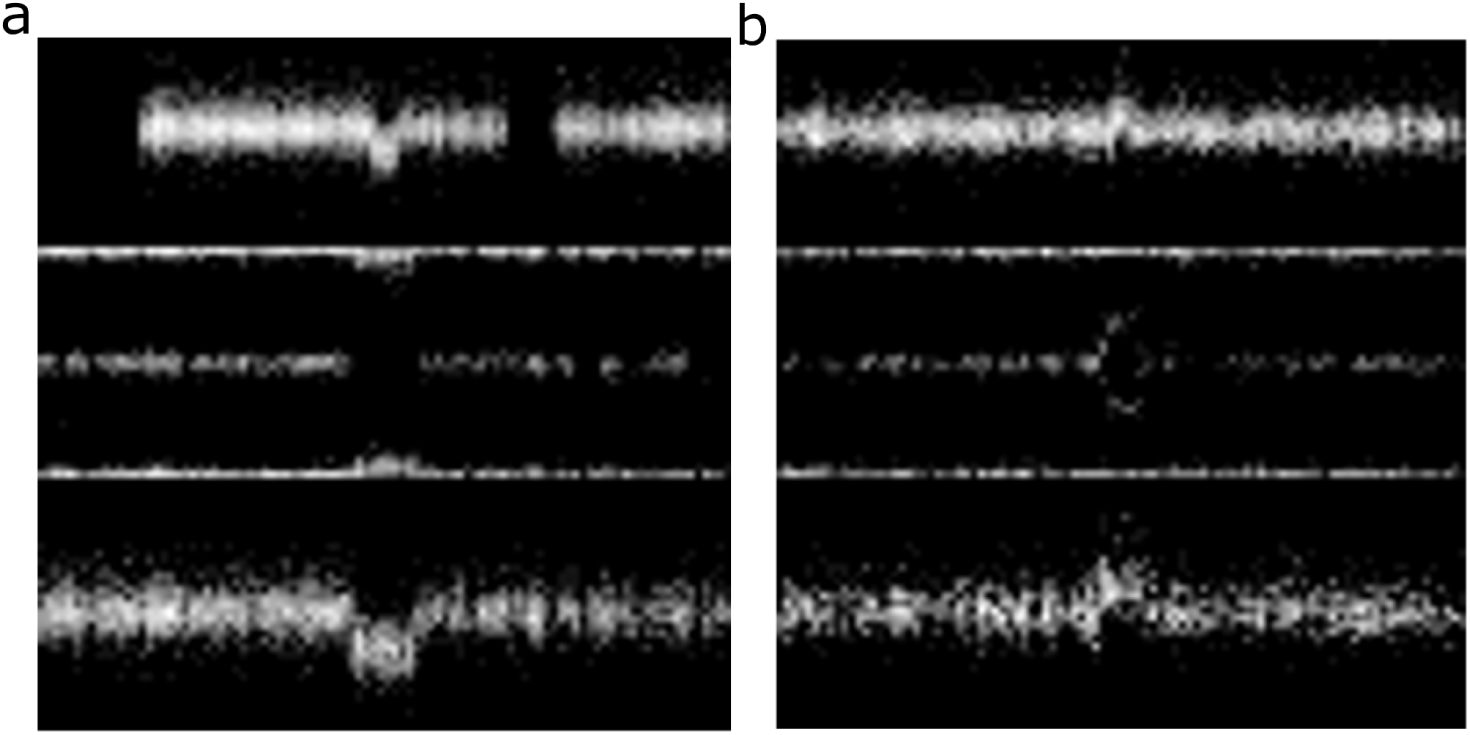
Example of a deletion and a duplication in the 96×96 pixels format for the model.

**Supplementary figure 3.**
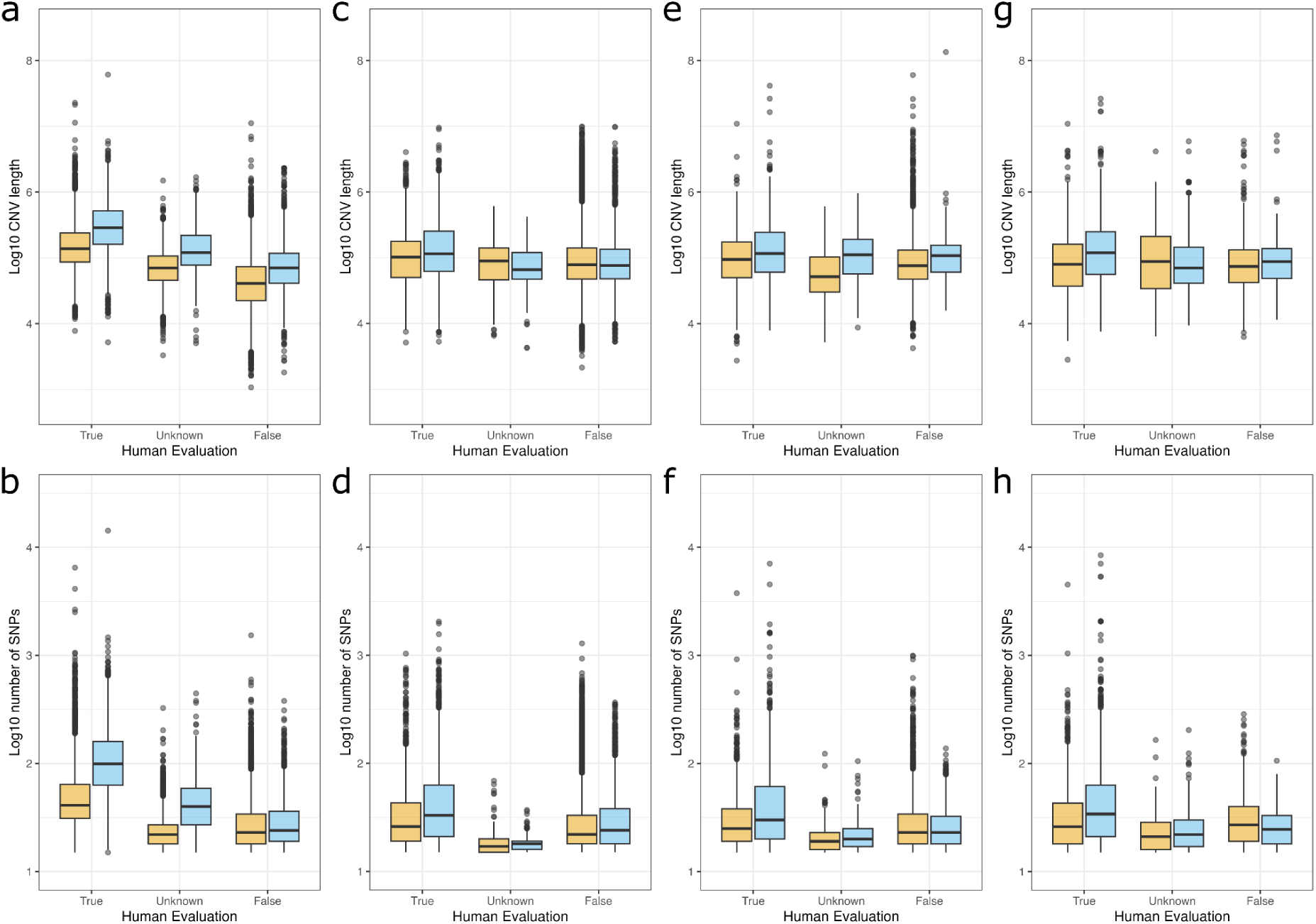
Log 10 of length (top) and number SNP makers (bottom) in the true, false and unclear CNVs. Deletions in yellow, duplications in blue. a-b) UKB (15,000 samples), c-d) deCODE (6,000 samples), e-f) 1kG HD (1,500 samples) and g-h) 1kG GW (1,500 samples). The y-axis range is kept consistent across each row whenever possible, provided it does not compromise the visualization.

**Supplementary figure 4.**
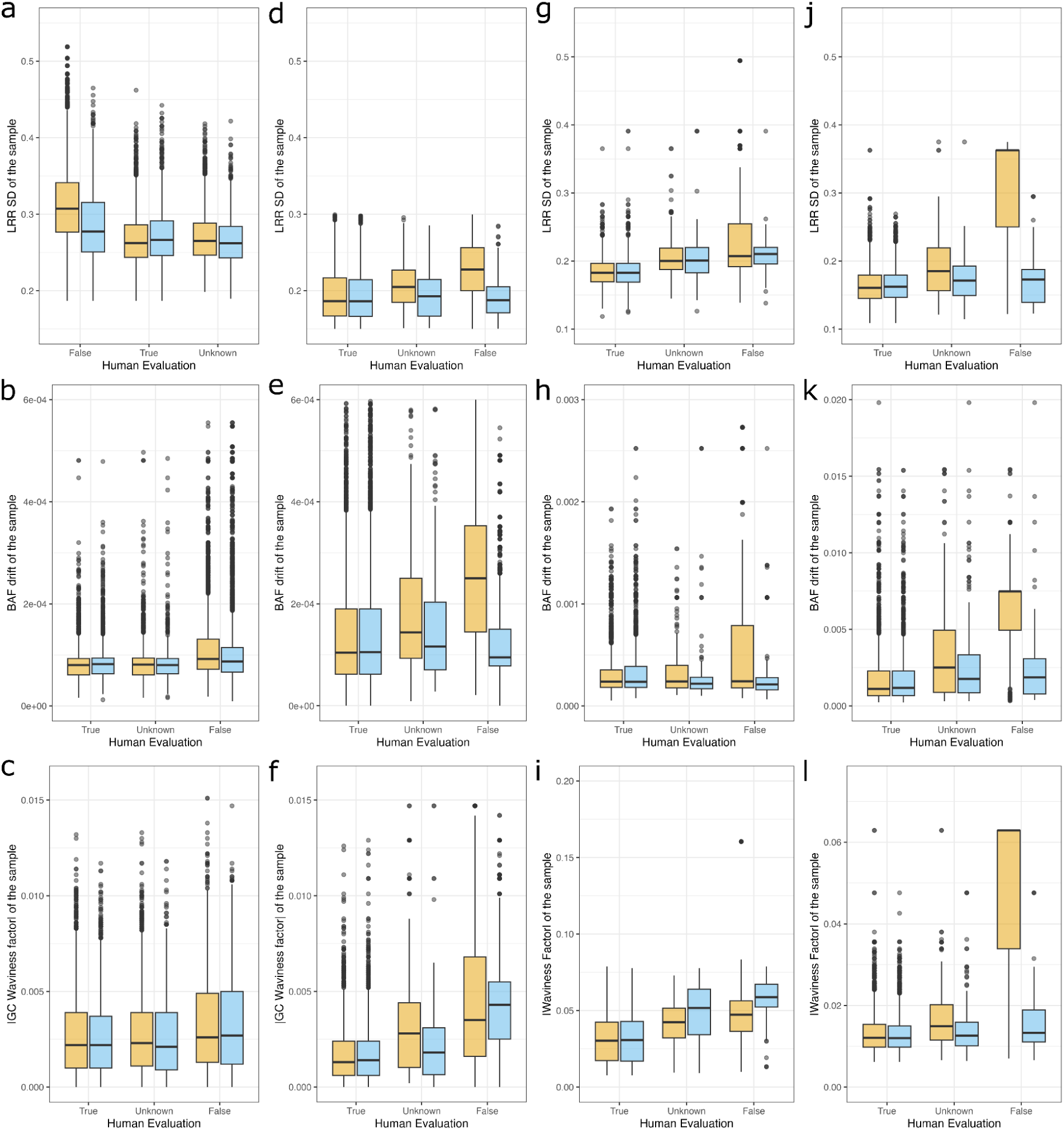
LRR SD (top), BAF drift (centre), and GCWF (bottom) distributions in the true, false and unclear CNVs. Deletions in yellow, duplications in blue. Each CNV inherits the QC measure of the sample. a-c) UKB (15,000 samples), d-f) deCODE (6,000 samples), g-i) 1kG HD (1,500 samples) and j-l) 1kG GW (1,500 samples). The y-axis range is kept consistent across each row whenever possible, provided it does not compromise the visualization.

**Supplementary figure 5.**
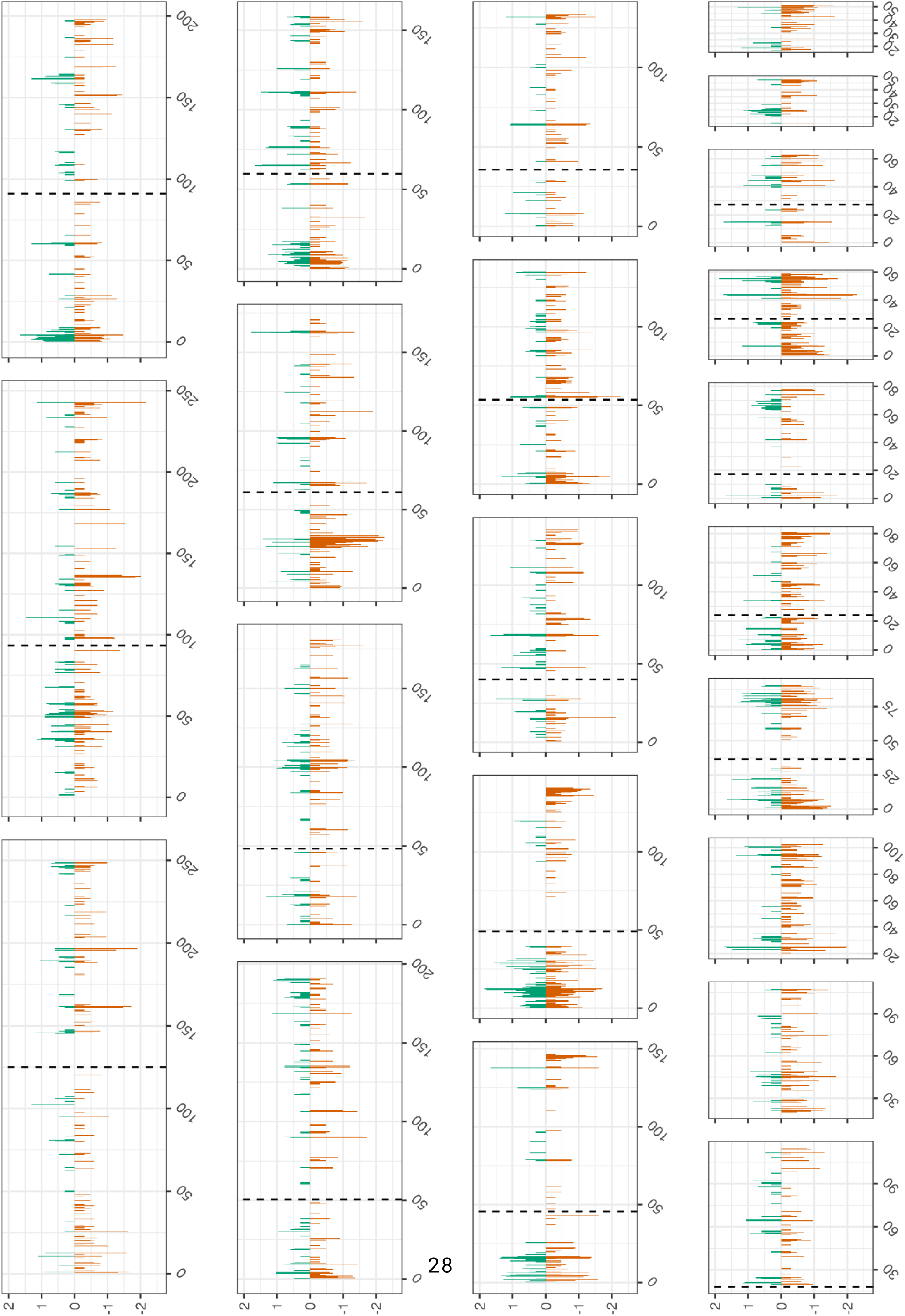
True (green) and false (orange) positive CNV calls genomic distribution for UKB deletions. Each bar represents the number of CNV in the 500kbp window on the log10 scale.

**Supplementary figure 6.**
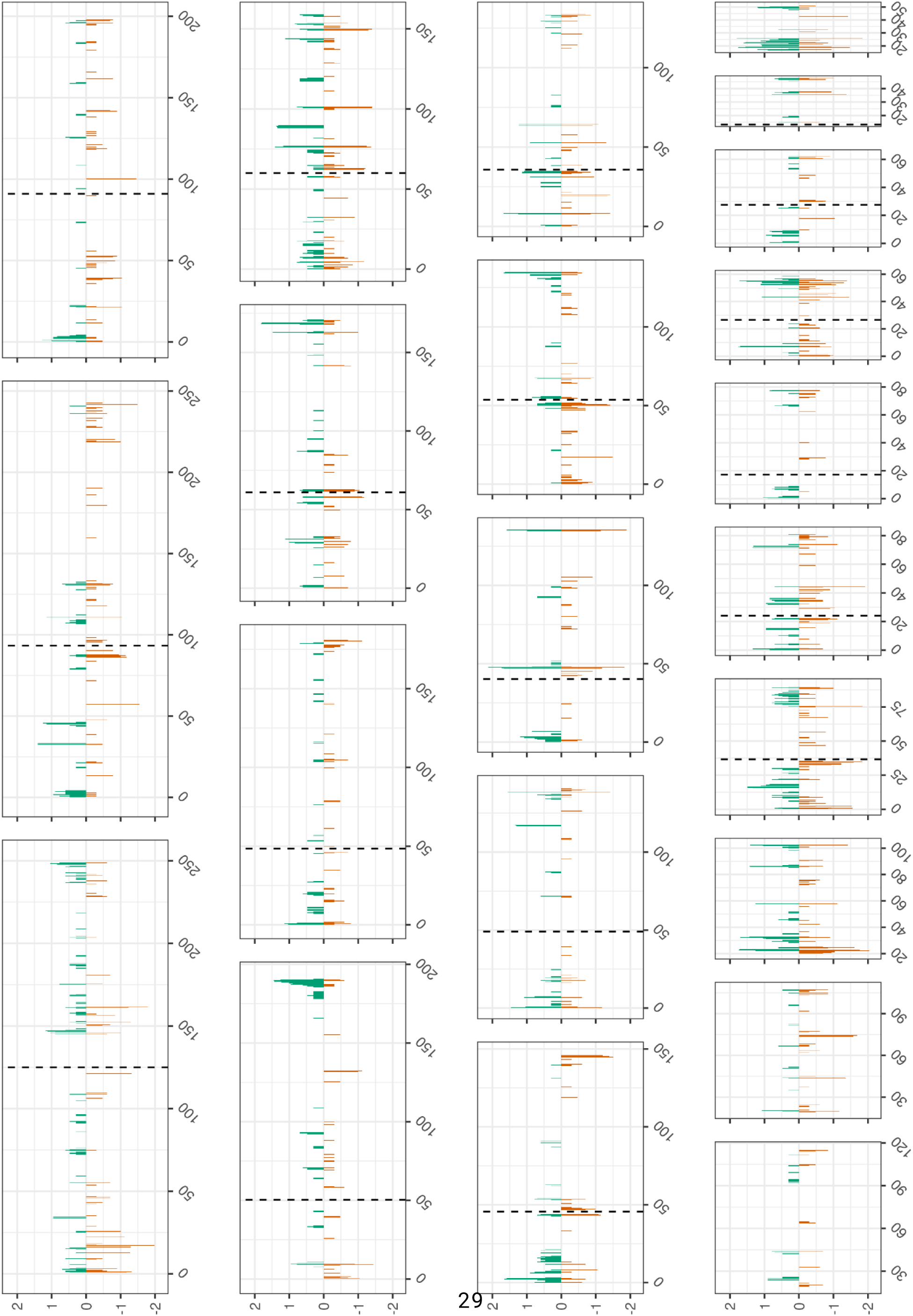
True (green) and false (orange) positive CNV calls genomic distribution for UKB duplications. Each bar represents the number of CNV in the 500kbp window on the log10 scale.

**Supplementary figure 7.**
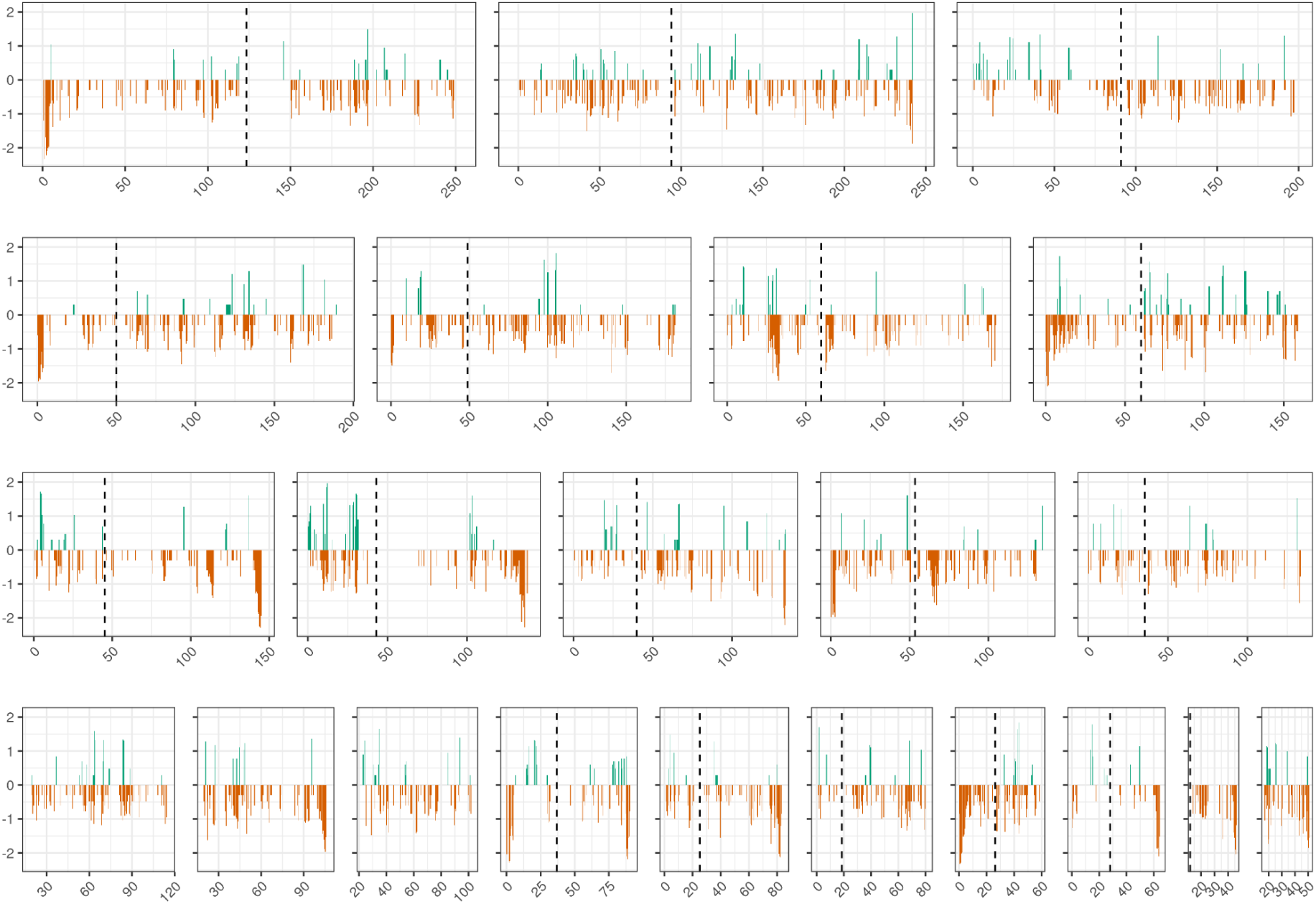
True (green) and false (orange) positive CNV calls genomic distribution for deCODE deletions. Each bar represents the number of CNV in the 500kbp window on the log10 scale.

**Supplementary figure 8.**
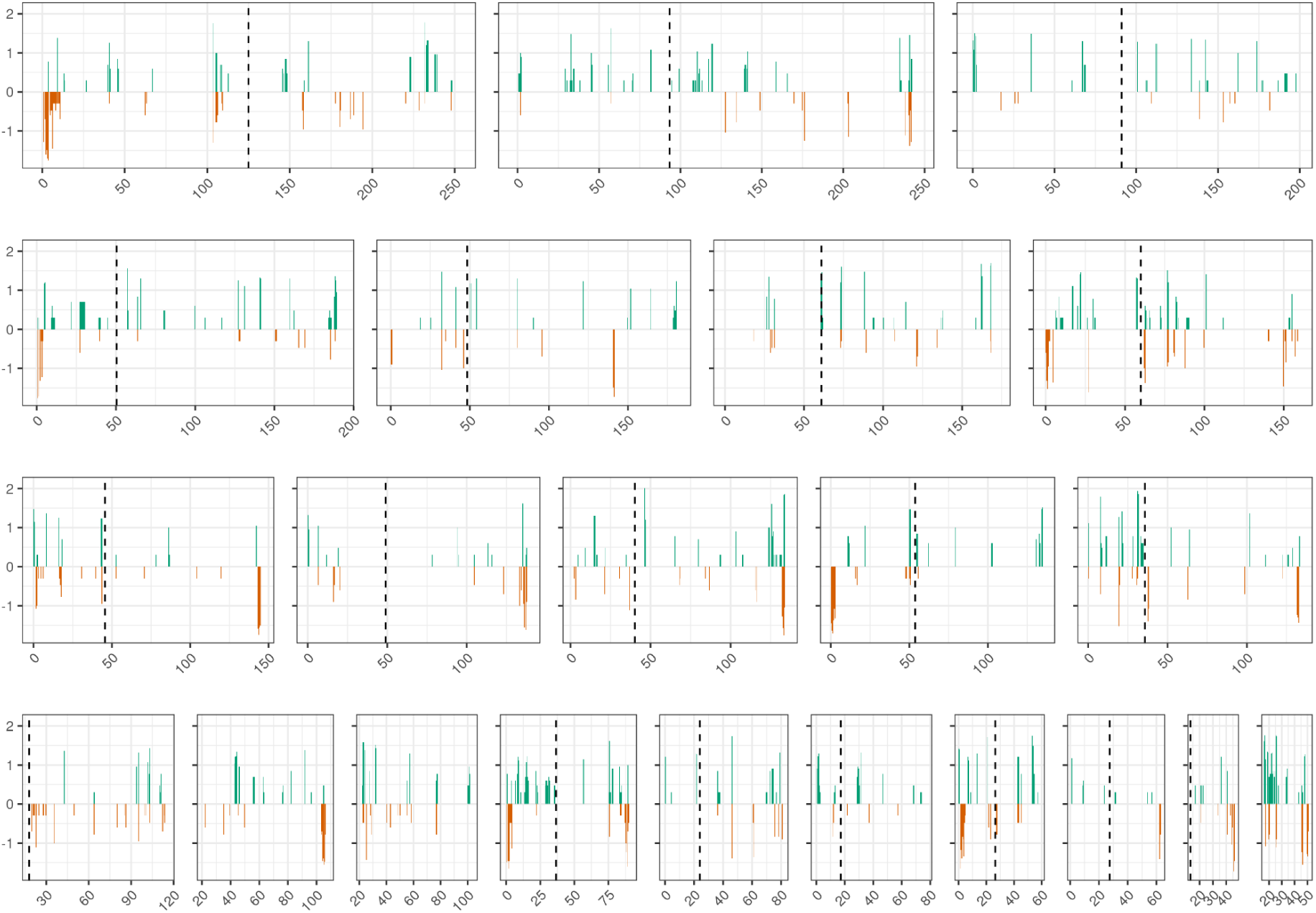
True (green) and false (orange) positive CNV calls genomic distribution for deCODE duplications. Each bar represents the number of CNV in the 500kbp window on the log10 scale.

**Supplementary figure 9.**
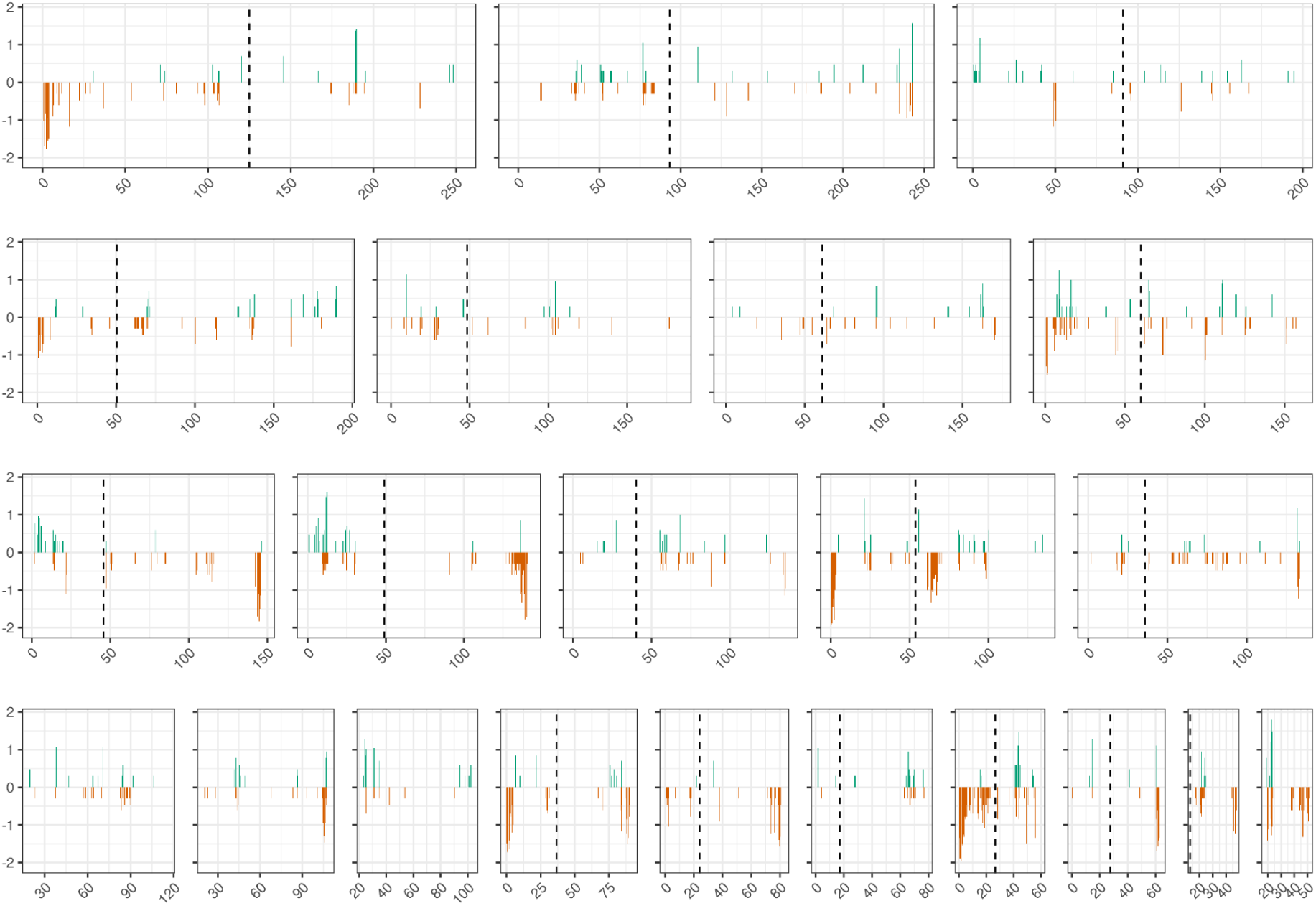
True (green) and false (orange) positive CNV calls genomic distribution for 1kG HD deletions. Each bar represents the number of CNV in the 500kbp window.

**Supplementary figure 10.**
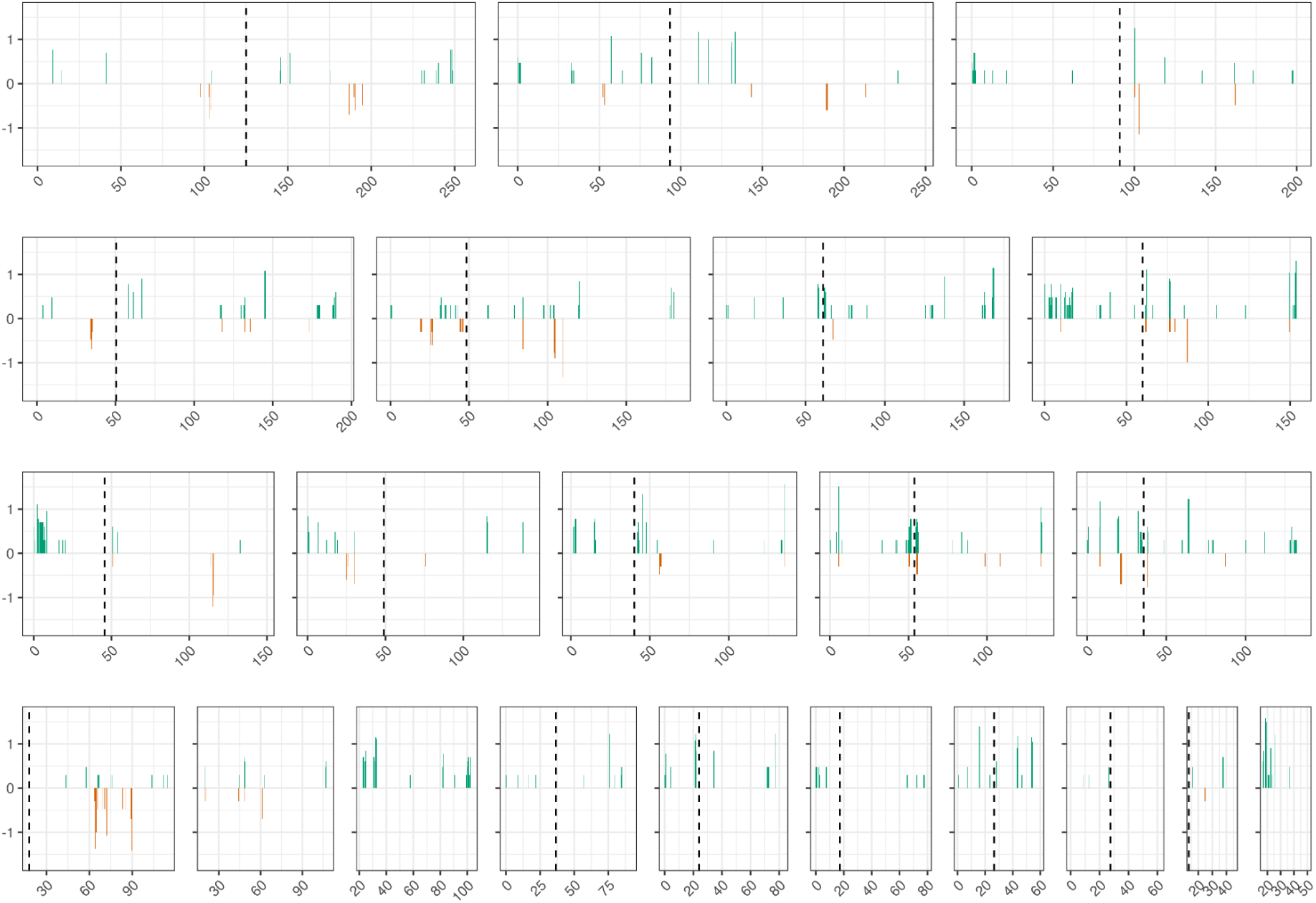
True (green) and false (orange) positive CNV calls genomic distribution for 1kG HD duplications. Each bar represents the number of CNV in the 500kbp window.

**Supplementary figure 11.**
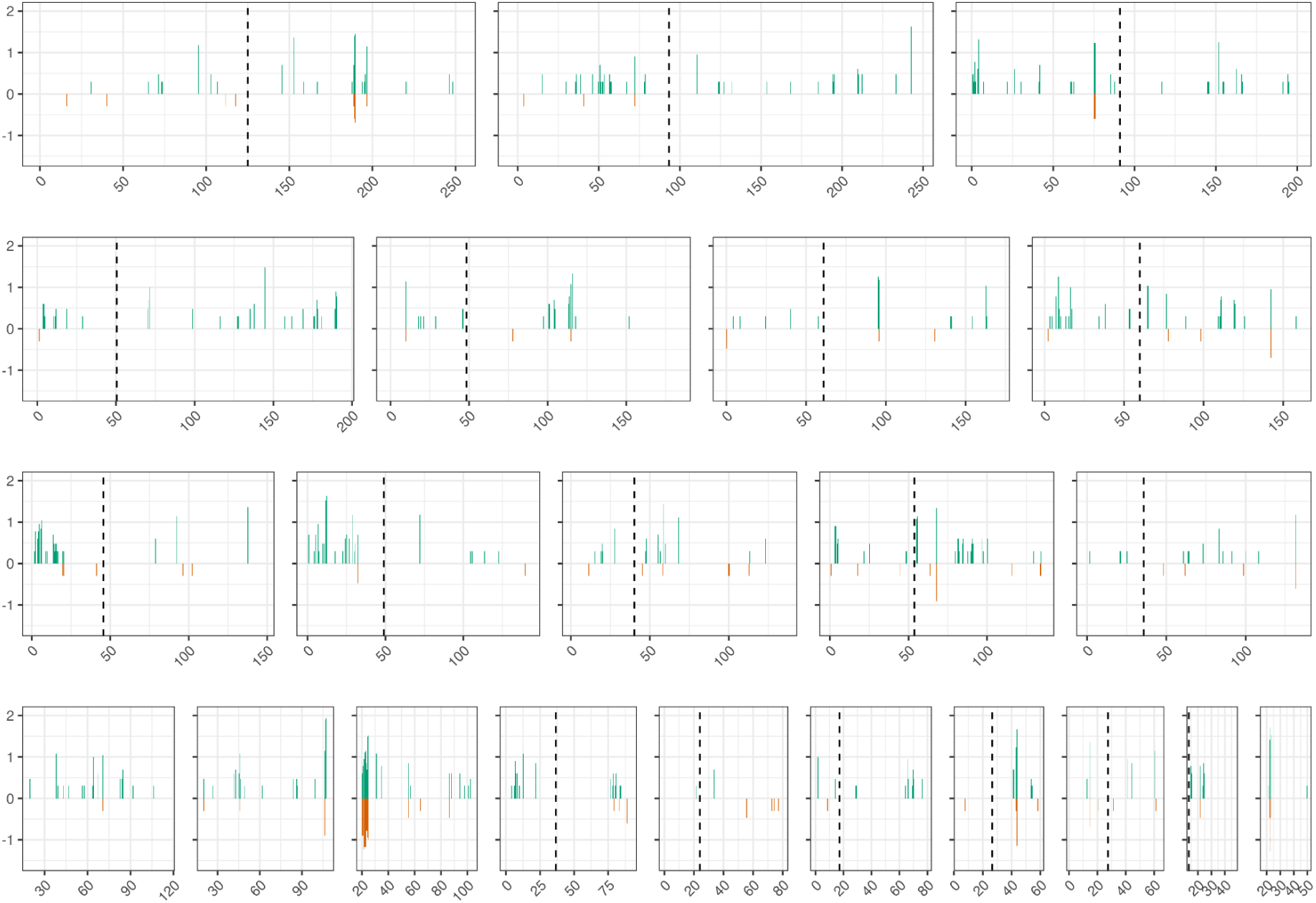
True (green) and false (orange) positive CNV calls genomic distribution for 1kG GW deletions. Each bar represents the number of CNV in the 500kbp window.

**Supplementary figure 12.**
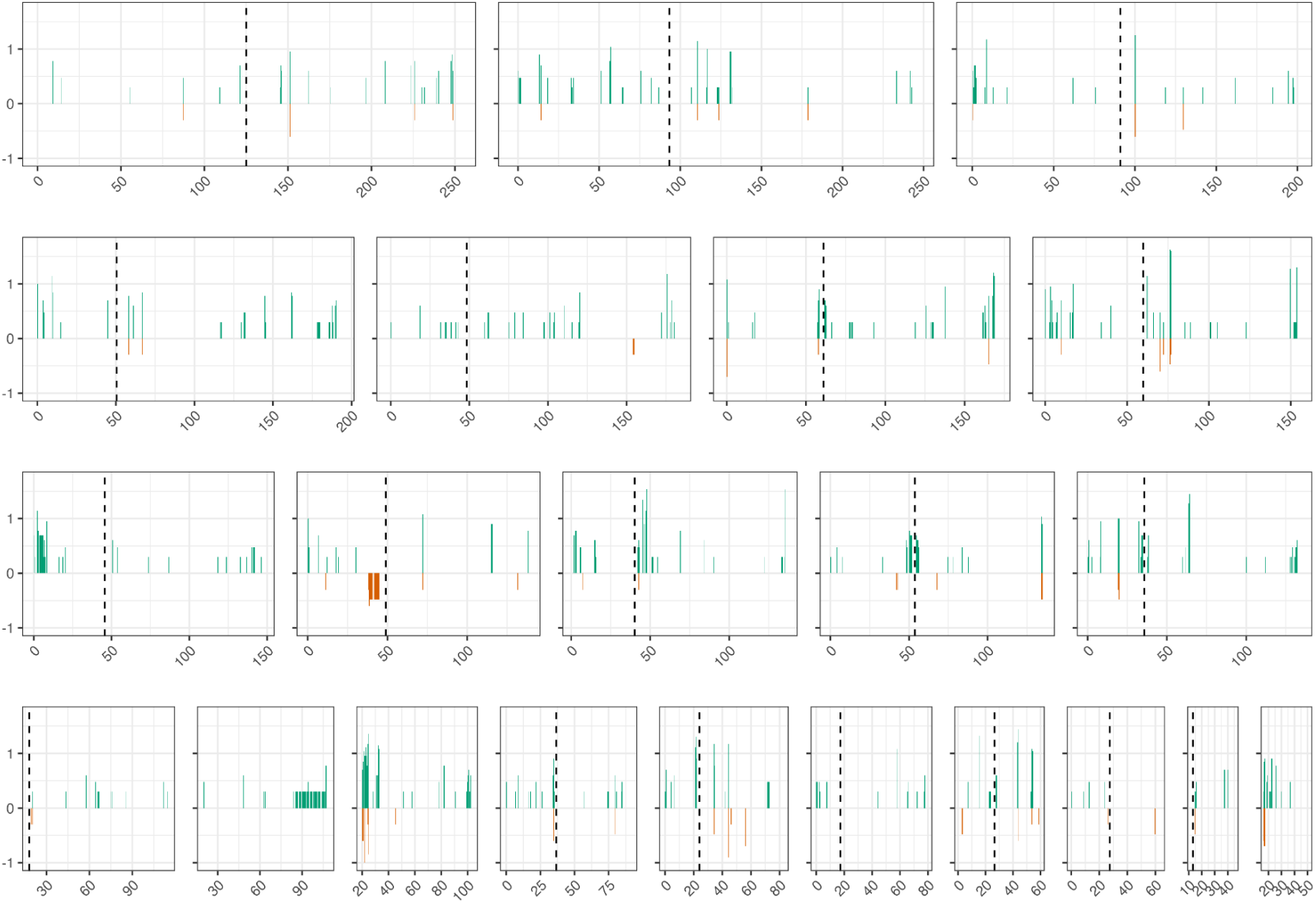
True (green) and false (orange) positive CNV calls genomic distribution for 1kG GW duplications. Each bar represents the number of CNV in the 500kbp window.

